# Crowder-specific modulation of hepatitis C virus NS3/4A protease activity and local structural dynamics

**DOI:** 10.64898/2026.02.27.708426

**Authors:** Małgorzata Lobka, Joanna Trylska

## Abstract

Macromolecular crowding modulates enzyme behavior in crowder- and protein-specific ways, yet its impact on viral proteases, which are often key therapeutic targets, remains unclear. Here, we investigated the hepatitis C virus NS3/4A protease under increasing concentrations of polyethylene glycols (PEGs), ficoll, dextran, and lysozyme using a fluorescence-based activity assay and intrinsic tryptophan fluorescence. PEGs reduced catalytic activity while leaving substrate binding largely unaffected or moderately enhanced. These effects were accompanied by a moderate tryptophan fluorescence spectral narrowing, consistent with reduced heterogeneity in local conformational environments. In contrast, ficoll enhanced catalytic efficiency despite stronger fluorescence quenching, indicating local structural changes that favored catalysis. Dextran and lysozyme inhibited protease activity through distinct kinetic patterns, likely reflecting differences in their size, shape, and chemical properties. Thermal analysis revealed crowder-specific local structural changes in NS3/4A without global unfolding up to 65°C, with differences in local stability and flexibility corroborating the observed kinetic effects. These findings demonstrate that macromolecular crowding modulates NS3/4A catalysis through crowder-specific effects on local structure.

## 1 Introduction

Although approximately 85% of a typical eukaryotic cell is water [1], the remaining volume is densely packed with macromolecules such as proteins, lipids, polysaccharides, and nucleic acids [2]. This crowding, present in both cytoplasm and membranes, limits molecular freedom through steric exclusion. The excluded-volume effect favors compact conformations and entropically stabilizes associations by minimizing surface area [3, 4]. In contrast, the chemical nature of macromolecules gives rise to nonspecific interactions that can favor extended conformations by maximizing contact area [5]. Together, steric and soft interactions modulate the thermodynamics and kinetics of diffusion, folding, ligand binding, and stability [4–6]. These effects are particularly relevant for enzymes, whose activity depends on the structural flexibility required for substrate binding and catalysis [7–9]. For viral enzymes, folding determines not only catalytic competence but also engagement with host factors regulating replication and immune defense [10]. Understanding how the environment modulates viral enzyme function may therefore provide mechanistic insights and support therapeutic development. However, predicting how folding and kinetics are altered under crowded conditions remains challenging, as crowder size, shape, and chemical nature—together with enzyme-specific properties—can modulate or override excluded-volume effects [5, 11–14].

The hepatitis C virus (HCV), a major cause of chronic liver disease and hepatocellular carcinoma [15], encodes the NS3/4A protease, which cleaves the viral polyprotein to release mature proteins essential for replication [16]. NS3 provides the N-terminal serine protease domain (aa 1–180) and, for in vivo catalysis, uses a 54-amino-acid cofactor, NS4A. However, only its central hydrophobic segment (aa 21–32), embedded in the NS3 core, is sufficient for in vitro activity [17]. This segment is hereafter referred to as NS4A. The protease adopts a chymotrypsin-like fold stabilized by zinc, with catalytic organization dependent on NS4A binding (see **Figure 1A**). Zinc removal disrupts the native fold and disorganizes the catalytic triad [18, 19]. NS4A binding stabilizes the heterodimer and organizes the triad, particularly by positioning His57 for proteolysis [20, 21]. In the absence of NS4A, catalytic residues remain mobile and only transiently hydrogen-bonded [22]. Beyond replication, the NS3/4A protease also cleaves the innate immune adaptors Cardif and Trif [23], contributing to hepatitis C pathogenesis. This dual function makes NS3/4A a validated therapeutic target. Three generations of substrate-mimicking inhibitors have been approved by the U.S. Food and Drug Administration [24], and compounds targeting non-catalytic regions have been identified as well [25]. However, resistance mutations limit long-term efficacy [26]. Since intracellular conditions can modulate NS3/4A function, studying this enzyme under crowding may reveal features absent in dilute solutions. Such insights could support the design of more resilient antiviral strategies. Methods for studying NS3/4A in vivo are under active development [27– 29]. However, few studies have examined NS3/4A under conditions mimicking intracellular complexity, either in silico [30–32] or in vitro [33, 34].

**Fig. 1.**
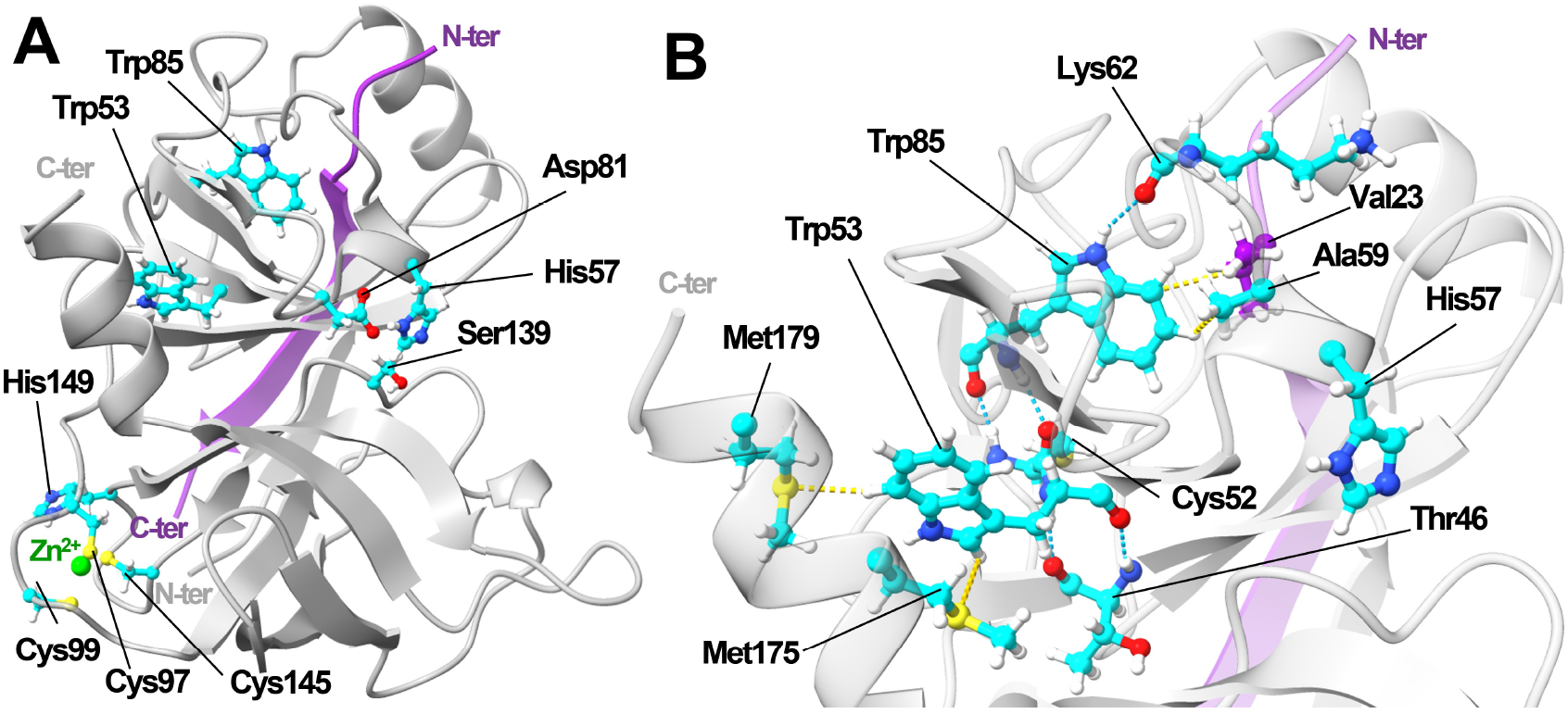
Structure of the NS3/4A protease functional complex. (A) Ribbon model of NS3 protease (grey) with NS4A (purple), visualized in ChimeraX [35], based on previous simulations [34] and crystal structure PDB ID: 4JMY [36]. Catalytic triad (His57, Asp81, Ser139), zinc-binding site (Cys97, Cys99, Cys145, His149; zinc ion in green), and tryptophan residues are highlighted in cyan. (B) Local environment of tryptophans with selected neighbors (for full contact lists, see **Table S1, Table S2**). Blue dashed lines indicate hydrogen bonds with distances of 2.8–3.1 Å between donor and acceptor; yellow dashed lines show selected hydrophobic contacts of 2.8–3.7 Å; Val23 (from NS4A) is shown in purple.

To mimic crowding in vitro, synthetic non-ionic polymers such as polyethylene glycol (PEG; a flexible polyether), ficoll (a highly branched sucrose-based copolymer), or dextran (a less branched polymer of glucose) are widely used. These polymers can influence protein behavior not only through excluded volume but also via hydrogen bonding, hydrophobic contacts, electrostatic forces, and van der Waals interactions, leading to diverse effects [5]. PEG often reduces enzyme activity, as observed for oxidoreductases and hydrolases [37–39], though activity enhancements have also been reported at low concentrations or with specific enzyme–substrate pairs [40]. Ficoll more frequently enhances activity, as seen for dehydrogenases and transferases [11, 41], though inhibition has been observed at high concentrations or in uncompetitive-like systems [42, 43]. Dextran most often reduces kinetic rates, especially above the chain overlap concentration [44–47], although increased activity at low concentrations has also been reported [11]. These crowders also affect protein folding. PEG can destabilize some proteins [48, 49], but in others it favors compact states [50]. Ficoll generally stabilizes folding [51, 52], though destabilization of partially folded states has also been described [53]. Dextran tends to stabilize globular proteins [47, 54]; however, its effect is not universally stabilizing and may even induce local destabilization or conformational changes [55].

Given this variability, it remains unclear how crowding influences NS3/4A function. Our previous work showed opposing effects on catalytic activity and inhibition [34]: PEG 600, PEG 6000, and bovine serum albumin (BSA) reduced activity, whereas Ficoll 400 enhanced it, with the initial reaction rate changing in a concentration-dependent manner. Ficoll also increased the inhibition constant for the antiviral drug telaprevir. Molecular dynamics (MD) simulations suggested that PEG slows substrate diffusion and increases substrate configurational entropy [31]. Further simulations indicated that ficoll stabilizes extended substrate conformations and promotes productive substrate encounters near the NS3/4A catalytic triad [32]. Following these contrasting observations, we examined the effects of PEG and ficoll of different molecular weights, as well as Dextran 70 and lysozyme, not only on NS3/4A activity but also on the protease’s local structure and thermal stability.

## 2 Results and Discussion

### 2.1 NS3/4A-mediated substrate hydrolysis

The NS3/4A kinetics were analyzed under dilute and crowded conditions by determining initial rate (*v*_0_), maximal rate (*v*_max_), Michaelis constant (*K*_M_), turnover number (*k*_cat_), and catalytic efficiency (*k*_cat_ /*K*_M_). *K*_M_ represents the substrate concentration at half-maximal velocity and reflects both substrate affinity and turnover [56]. *k*_cat_ quantifies how many substrates are converted per enzyme per unit time and reflects the rate-limiting step along the reaction pathway. The ratio *k*_cat_ /*K*_M_ describes catalytic performance under low substrate concentrations ([S] ≪ *K*_M_) [56]. Substrate-concentration-dependent changes in *v*_0_ are shown in **Figure 2A**, and the derived kinetic parameters are summarized in **Table 1 and Figure 3**.

**Table 1.** Kinetic parameters of NS3/4A in dilute and crowded conditions. *v*_max_, *K*_M_, and *k*_cat_ were derived from Michaelis– Menten fits (**Figure 2A**) and are reported as mean±SEM. SEMs of *k*_cat_/*K*_M_ were propagated from fitted parameters using standard error propagation including *K*_M_ and *k*_cat_ covariance. For each condition, excluded volume fractions (ϕ) and effective substrate concentration factors ([S]_eff_/[S]) were calculated. Statistical significance of *v*_max_, *K*_M_, and *k*_cat_ relative to dilute conditions is shown in **Figure 3**.

| Crowder [g/L] | | $\phi$ | $[S]_{\text{eff}}/[S]$ | $v_{\max}$ [$\mu\text{M/s}$] | $K_M$ [$\mu\text{M}$] | $k_{\text{cat}}$ [$\text{s}^{-1}$] | $k_{\text{cat}}/K_M$ [$\mu\text{M}^{-1}\text{s}^{-1}$] |
| --- | --- | --- | --- | --- | --- | --- | --- |
| – | 0<sup>a</sup> | n/a | n/a | $7.8 \pm 0.1$ | $19 \pm 1$ | $3,900 \pm 100$ | $205 \pm 5$ |
| PEG 2000 | 50 | 0.1 | 1.2 | $5.9 \pm 0.1$ | $17 \pm 1$ | $2,800 \pm 100$ | $165 \pm 5$ |
| | 100 | 0.3 | 1.4 | $4.8 \pm 0.3$ | $16 \pm 2$ | $2,300 \pm 100$ | $144 \pm 12$ |
| | 200 | 0.5 | 2.1 | $4.3 \pm 0.4$ | $14 \pm 2$ | $2,200 \pm 200$ | $157 \pm 11$ |
| PEG 4000 | 50 | 0.2 | 1.2 | $6.5 \pm 0.1$ | $12 \pm 1$ | $3,300 \pm 100$ | $275 \pm 16$ |
| | 100 | 0.3 | 1.5 | $5.8 \pm 0.2$ | $12 \pm 1$ | $2,900 \pm 100$ | $242 \pm 13$ |
| | 200 | 0.6 | 2.8 | $4.1 \pm 0.2$ | $11 \pm 1$ | $2,100 \pm 100$ | $191 \pm 10$ |
| Ficoll 70 | 50 | 0.2 | 1.2 | $8.9 \pm 0.2$ | $19 \pm 1$ | $4,400 \pm 100$ | $232 \pm 7$ |
| | 100 | 0.4 | 1.6 | $13.4 \pm 0.6$ | $29 \pm 2$ | $6,700 \pm 300$ | $231 \pm 6$ |
| | 200 | 0.8 | 4.0 | $19.0 \pm 0.5$ | $25 \pm 1$ | $9,500 \pm 300$ | $380 \pm 5$ |
| Dextran 70 | 50 | 0.5 | 2.0 | $6.1 \pm 0.2$ | $14 \pm 1$ | $3,000 \pm 100$ | $214 \pm 9$ |
| | 100 | $\approx 1.0$ | 67.1 | $5.4 \pm 0.3$ | $15 \pm 2$ | $2,700 \pm 100$ | $180 \pm 18$ |
| | 200 | n/a | n/a | $5.1 \pm 0.1$ | $3 \pm 1$ | $2,600 \pm 100$ | $867 \pm 263$ |
| Lysozyme | 15 | $< 0.1$ | 1.0 | $1.1 \pm 0.1$ | $3 \pm 1$ | $600 \pm 100$ | $200 \pm 42$ |
| | 25 | $< 0.1$ | 1.0 | $0.8 \pm 0.1$ | $2 \pm 1$ | $400 \pm 100$ | $200 \pm 63$ |
| | 50 | 0.1 | 1.1 | $0.4 \pm 0.1$ | $1 \pm 1$ | $200 \pm 100$ | $200 \pm 125$ |

<sup>a</sup>Previously reported kinetic parameters (mean ± SEM; n=3) in dilute conditions provided for comparison: $v_{\max} = 3.4 \pm 0.1$ $\mu\text{M/s}$ ; $K_M = 7 \pm 1$ $\mu\text{M}$ ; $k_{\text{cat}} = 1,700$ $\text{s}^{-1}$ [34].

**Fig. 2.**
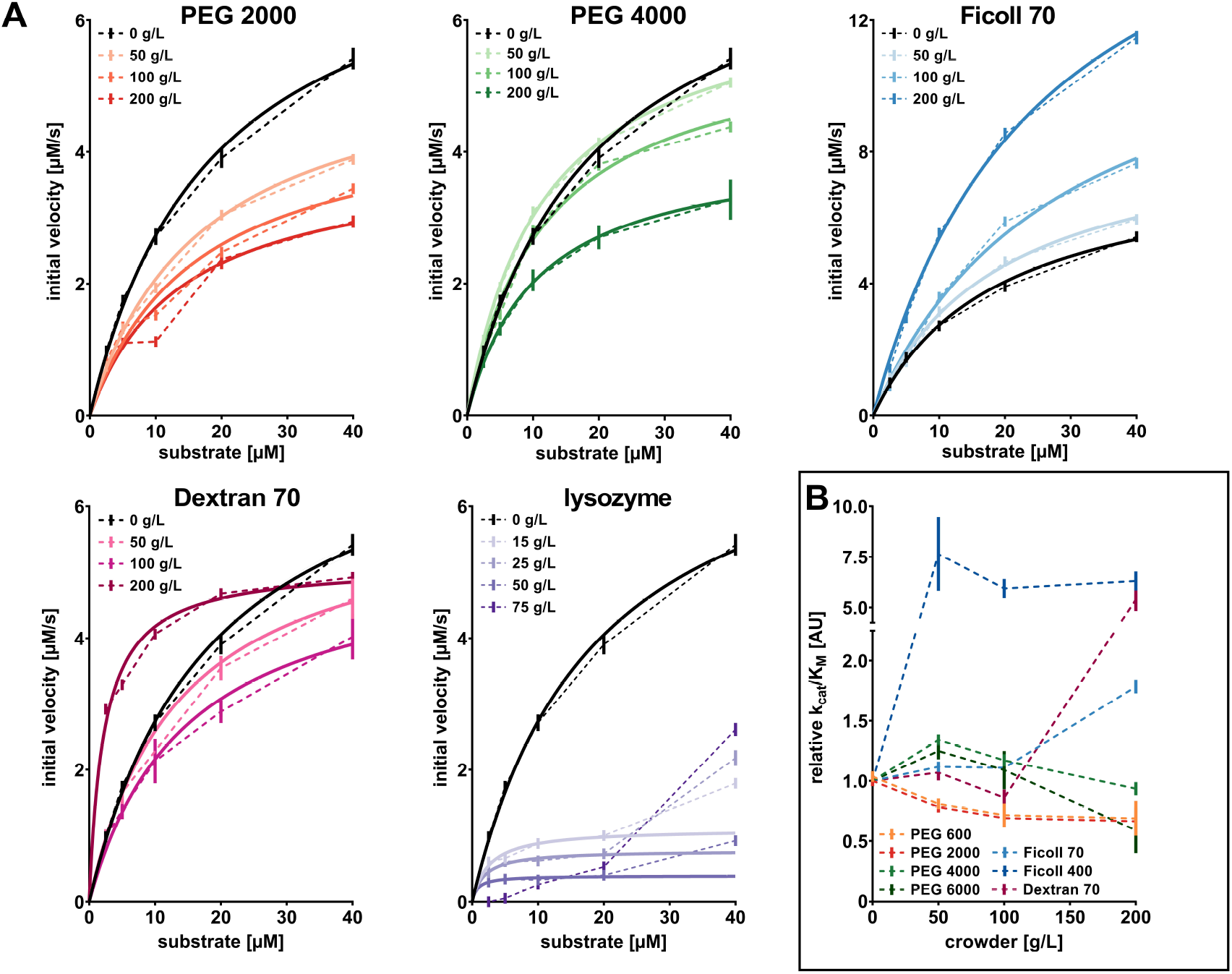
NS3/4A-mediated hydrolysis kinetics under dilute and crowded conditions. (A) Initial velocities were determined using 0-40 µM substrate and 0.05 µg/mL enzyme in dilute conditions, 50-200 g/L polymer crowders, and 15-75 g/L lysozyme. Data points represent mean±SEM (n = 3, each performed in duplicate) and were fitted to the Michaelis-Menten equation at the 0-40 µM range for polymer crowders and 0-20 µM for lysozyme (solid lines). Statistical differences between reaction rates at 40 µM substrate relative to dilute conditions were assessed by two-way ANOVA and were significant for all crowding agents and concentrations (p<0.0001). (B) Comparison of relative catalytic efficiencies (*k*_cat_/*K*_M_), shown as mean ± SEM (n = 3, each in duplicate). Data for PEG 600, PEG 6000, and Ficoll 400 were taken from [34].

**Fig. 3.**
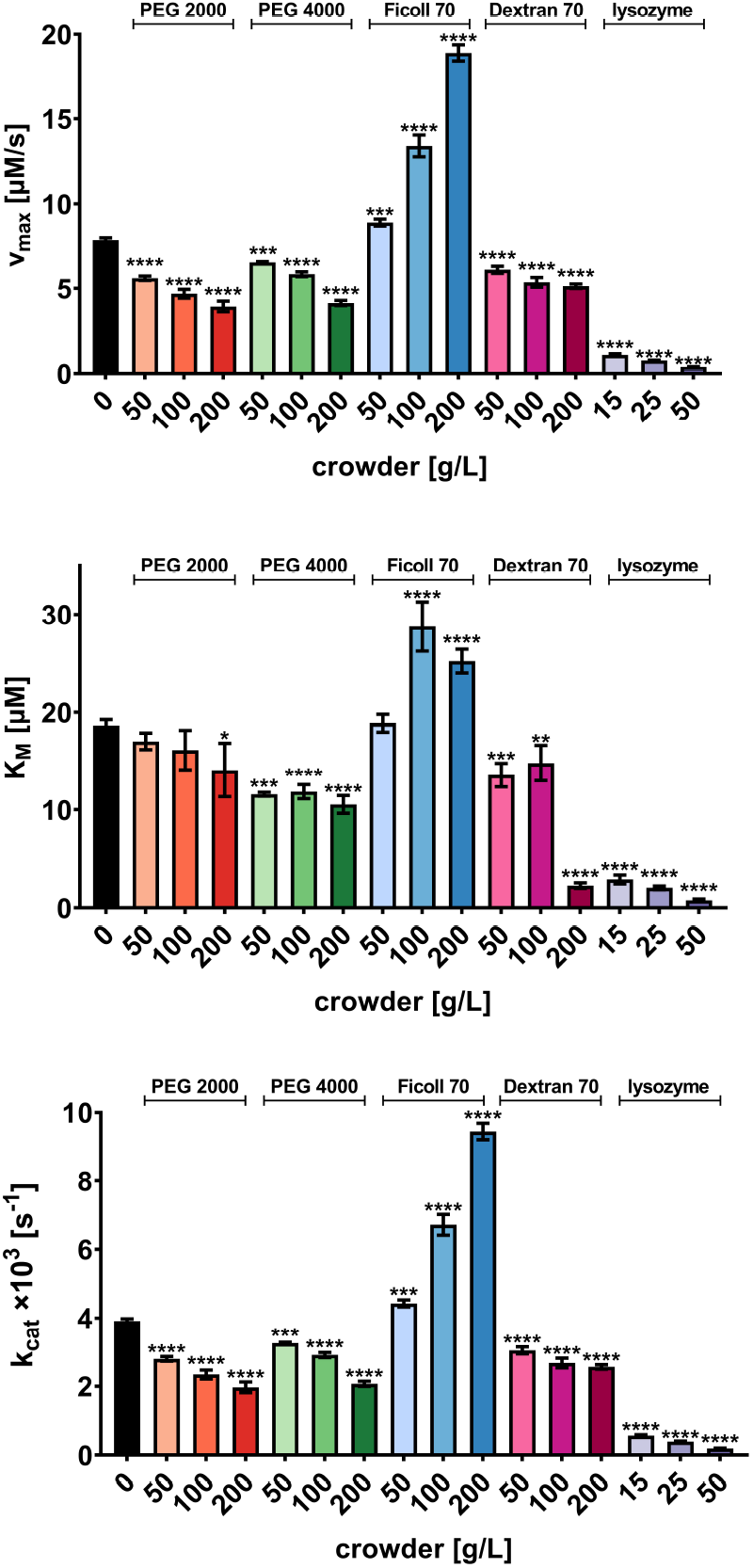
Kinetic parameters of NS3/4A in dilute and crowded conditions. Maximal velocity *v*_max_, Michaelis constant *K*_M_, and turnover number *k*_cat_ were derived from Michaelis–Menten fits (**Figure 2A**) and are reported as mean ± SEM (n = 3, each in duplicate). Statistical significance of *v*_max_, *K*_M_, and *k*_cat_ relative to dilute conditions was evaluated by two-way ANOVA and is denoted as ****p < 0.0001, ***p < 0.001, **p < 0.01, and *p < 0.05; otherwise, differences were not statistically significant.

Under dilute conditions, NS3/4A catalytic efficiency at low substrate concentrations corroborated our previous report [34], but both *k*_cat_ and *K*_M_ were approximately 2.3–2.7-fold higher (**Table 1**). This pattern (higher *k*_cat_ and *K*_M_ but stable *k*_cat_ /*K*_M_) suggests that the rate-limiting transition state was unchanged but the energy of a preceding intermediate (e.g., the enzyme-substrate complex) was altered [56]. This shift in the enzyme energy landscape likely resulted from a revised enzyme-handling protocol; instead of thawing NS3/4A directly at room temperature [34], the enzyme was preincubated on ice before equilibration at room temperature (see **Buffer and sample preparation** in **Materials and methods**), as is commonly done to prevent thermal shock [57]. However, such preincubation may promote local unfolding through ice recrystallization [58, 59], potentially explaining the higher turnover number and the concurrent increase in *K*_M_ . A similar trade-off has been reported, e.g., for adenylate kinase following cold-induced conformational transitions [60].

The addition of PEG 2000 or PEG 4000 decreased both *k*_cat_ and *v*_max_, relative to dilute condition (**Table 1** and **Figure 2A**), reaching a 1.8–1.9-fold reduction at 200 g/L PEGs. The NS3/4A catalytic efficiency remained relatively stable across PEG concentrations, consistent with our observations for PEGs with other masses (PEG 600 and PEG 6000, **Figure 2B**). Overall, all tested PEGs reduced turnover in a concentration-dependent manner (**Figure 3**). Notably, *K*_M_ decreased modestly in 200 g/L PEG 2000 and at all PEG 4000 concentrations. These results indicate that PEG does not hinder substrate binding but instead slows the following catalytic steps [56]. This hypothesis aligns with our MD simulations showing that PEG slows substrate diffusion but increases substrate flexibility, and that PEG alters the NS3/4A conformational dynamics [32, 34]. Together, these factors likely reduce the probability of catalytically competent alignment of the enzyme-substrate complex. The modest decrease in *K*_M_ may partly reflect increased substrate availability due to excluded-volume effects, with effective substrate concentrations rising up to 2.1–2.8-fold in 200 g/L PEGs (**Table 1**). However, this increase did not compensate for the reduction in turnover, leading to decreased enzyme activity under saturating conditions. Similar PEG-dependent inhibition, arising from competing excluded-volume effects and soft interactions, has also been reported for other enzymes [37–39].

In contrast, with Ficoll 70, the kinetic parameters gradually increased relative to dilute conditions, reaching up to a 2.4-fold increase in reaction rate and 2.7-fold increase in turnover at 200 g/L (**Figure 2A** and **Figure 3**). At 100-200 g/L of Ficoll 70, *K*_M_ also increased, and at 200 g/L, the catalytic efficiency at sub-saturating conditions improved by 1.8-fold. A similar ficoll-induced enhancement of *k*_cat_ and *k*_cat_ /*K*_M_ was previously observed with Ficoll 400 (**Figure 2B**). The increase in both *k*_cat_ and *k*_cat_ /*K*_M_ is consistent with lowering of the rate-limiting transition-state energy barrier [56]. This effect can arise from destabilization of the enzyme-substrate complex before the chemical reaction, stabilization of a catalytically competent configuration, or narrowing of the energy barrier driven by fast molecular fluctuations [61]. Although these possibilities cannot be resolved with this kinetic assay, previous MD simulations of NS3/4A surrounded by tetra-sucrose molecules mimicking ficoll showed that this polymer forms soft interactions with the enzyme surface, stabilizes extended substrate conformations, and promotes productive encounters near the catalytic triad of NS3/4A [32]. This is consistent with the stronger stimulation of catalytic processing observed experimentally in this study. Despite improved turnover and catalytic efficiency, *K*_M_ increased at 100-200 g/L Ficoll 70, indicating that the enzyme required a higher substrate concentration to reach 1/2*v*_max_. However, the hydrophilic Ficoll 70 occupies up to 75% of the solution volume at 200 g/L, corresponding to nearly a 4-fold increase in effective substrate concentration (**Table 1**). Thus, the modest increase in *K*_M_ likely reflects the competing excluded volume effects and soft interactions between Ficoll, NS3/4A, and the FRET-substrate. Although Ficoll 400 appeared more effective than Ficoll 70 (**Figure S1**), slight differences in the applied protocols preclude a direct comparison. Similar ficoll-induced activity enhancement has also been reported for other enzymes [11, 41, 42].

In 50-200 g/L Dextran 70, a concentration-dependent decrease in kinetic rates was observed, with up to a 1.5-fold reduction in *v*_max_ and *k*_cat_ at 200 g/L relative to dilute conditions (**Table 1**). Catalytic efficiency remained stable up to 100 g/L, but increased by over 4-fold at 200 g/L due to an over 6-fold decrease in *K*_M_ (**Table 1, Figures 2B, 3**, and **Figure S1**). This abrupt shift likely reflects the polymer crossing its chain overlap concentration, which is 125 g/L for Dextran 70 in water [62]. Above this threshold, individual dextran polymer chains are no longer isolated; they begin to interpenetrate and entangle with one another, forming a transient meshwork, leading to abrupt kinetic alterations, as reported previously [62, 63]. In contrast, Ficoll 70 has a higher overlap concentration of 250 g/L [11], consistent with the absence of such a sharp transition in its kinetic effects at 50-200 g/L. The spherical approximation used for excluded-volume calculations also becomes invalid above 100 g/L Dextran 70 (**Table 1**), as the assumption of non-overlapping hard-sphere crowders no longer holds. Dextran-induced reduction in enzyme activity has been reported for other systems [44–46]. However, the opposite effects of ficoll and dextran on NS3/4A at 50-100 g/L are notable, as contrasting responses to polysaccharide crowders are relatively uncommon. Differences in magnitude have been reported previously [45, 64], whereas differences in the direction of kinetic changes have been described only in isolated cases, such as for yeast alcohol dehydrogenase [65]. These behaviors may arise from differences in polymer shape, flexibility, and viscosity effects, as well as from soft interactions with proteins. For example, the stronger inhibition of alkaline phosphatase activity by dextran relative to ficoll has been attributed to dextran’s extended, flexible structure, which introduces more physical barriers to enzyme-substrate encounters beyond viscosity effects alone [45].

As a protein crowder, we previously used BSA and determined the NS3/4A activity with up to 7.5 g/L of BSA [34]. Here, we used hen egg white lysozyme (HEWL) as an alternate protein crowder. We confirmed that the FRET-substrate is not cleaved by HEWL at its highest concentration tested (**Figure S2**), which allowed monitoring NS3/4A activity up to 75 g/L of lysozyme. However, at the highest HEWL concentrations, the NS3/4A kinetics deviated from the Michaelis–Menten model, with successful fitting only up to 20 µM FRET-substrate at 15-50 g/L of lysozyme (**Figure 2A**). The reaction rate continued to increase at 40 µM substrate without reaching a plateau, indicating non-hyperbolic behavior. A similar deviation was observed at 75 g/L across all substrate concentrations. Such deviation at [*S*] ≫ *K*_M_ has been reported and may reflect allosteric effects or other regulatory phenomena [66]. This aligns with reports that lysozyme shows more pronounced effects on enzyme kinetics, compared to polymer crowders or proteins with a weak crowding effect, such as BSA [65, 67]. Within the concentration range where Michaelis-Menten model fitting was still valid, lysozyme reduced NS3/4A activity substantially, with a nearly 20-fold decrease in *v*_max_ and *k*_cat_ at 50 g/L relative to dilute conditions (**Table 1**). Catalytic efficiency remained relatively stable due to a 19-fold reduction in *K*_M_, indicating that the rate-limiting transition state was not strongly affected [56]. Instead, lysozyme may have altered substrate binding dynamics or the enzyme’s active conformation, leading to slower turnover despite higher apparent affinity. Lysozyme’s surface contains positively charged lysines and arginines, which likely drive electrostatic and hydrogen-bonding interactions [68] with both the NS3/4A enzyme (**Figure S3**) and the negatively charged substrate. Such soft interactions are consistent with the modest excluded-volume fraction (up to 10% at 50 g/L; **Table 1**), indicating that lysozyme effects arise primarily from direct interactions rather than from steric crowding alone. A lysozyme-induced decrease in activity has also been reported for other enzymes [44, 69].

In summary, these results show that PEG and ficoll consistently produce opposite effects on NS3/4A catalysis across various molecular weights: PEG reduces activity, while ficoll enhances it. Although also a polysaccharide, dextran inhibits NS3/4A activity but increases catalytic efficiency above its chain overlap concentration. Lysozyme exerts stronger inhibition than the synthetic polymers, even at lower concentrations. To assess whether these changes involve structural alterations, we further examined the local tryptophan environment of NS3/4A under polymer crowding. Lysozyme was excluded from the fluorescence analysis due to its intrinsic tryptophan fluorescence, which would obscure signal attribution to NS3/4A.

### 2.2 Crowder-induced local structural changes near NS3/4A Trp residues

Protein dynamics are often dominated by minor conformational fluctuations, where the overall structural order remains intact but the distribution of accessible states changes [70]. To probe such local changes in NS3/4A under crowded conditions, we monitored its intrinsic tryptophan fluorescence. At 35°C, close to the temperature used in the kinetic assay, fluorescence was detectable at protein concentrations as low as 1 µg/mL. Samples containing the protein in buffer and crowder-containing solutions showed a significantly higher fluorescence yield than the background (**Figure 4A-B**), confirming a protein-specific signal. In dilute conditions, the emission maximum was 315 nm (**Figure 4A**), consistent with tryptophans located in a predominantly hydrophobic environment [71] and supported by our analysis of the NS3/4A structure (**Figure S3**). Both Trp residues have negligible solvent exposure (< 5 Å2). Trp53 is buried in the hydrophobic core between the NS3 β-sheet and C-terminal a-helix (**Figure 1A**) and forms contacts with Val51, Leu82, Val83, and Tyr75 (2.2–3.5 Å; **Table S1**), and hydrogen bonds with Thr46 (2.9 Å; **Figure 1B**). The nearby Met175 and Met179 side chains (2.8–3.8 Å) may influence fluorescence through weak sulfur-related quenching [72]. Trp53 also forms a small hydrophobic cavity positioned at the protein surface (**Figure S3**). Trp85 is positioned close to the NS3 surface loops and is stabilized by hydrogen bonds with Cys52 and Lys62 (2.8–3.1 Å; **Table S2**), and hydrophobic contacts with Leu64, Ile71, Val51, and Thr54 (1.8–3.4 Å). Although the reduced Cys52 is in proximity, it is not directly oriented toward the indole ring, suggesting low quenching probability [72]. Thus, both tryptophans report on tightly packed regions of NS3/4A and are well suited to detect local, rather than global, structural changes.

**Fig. 4.**
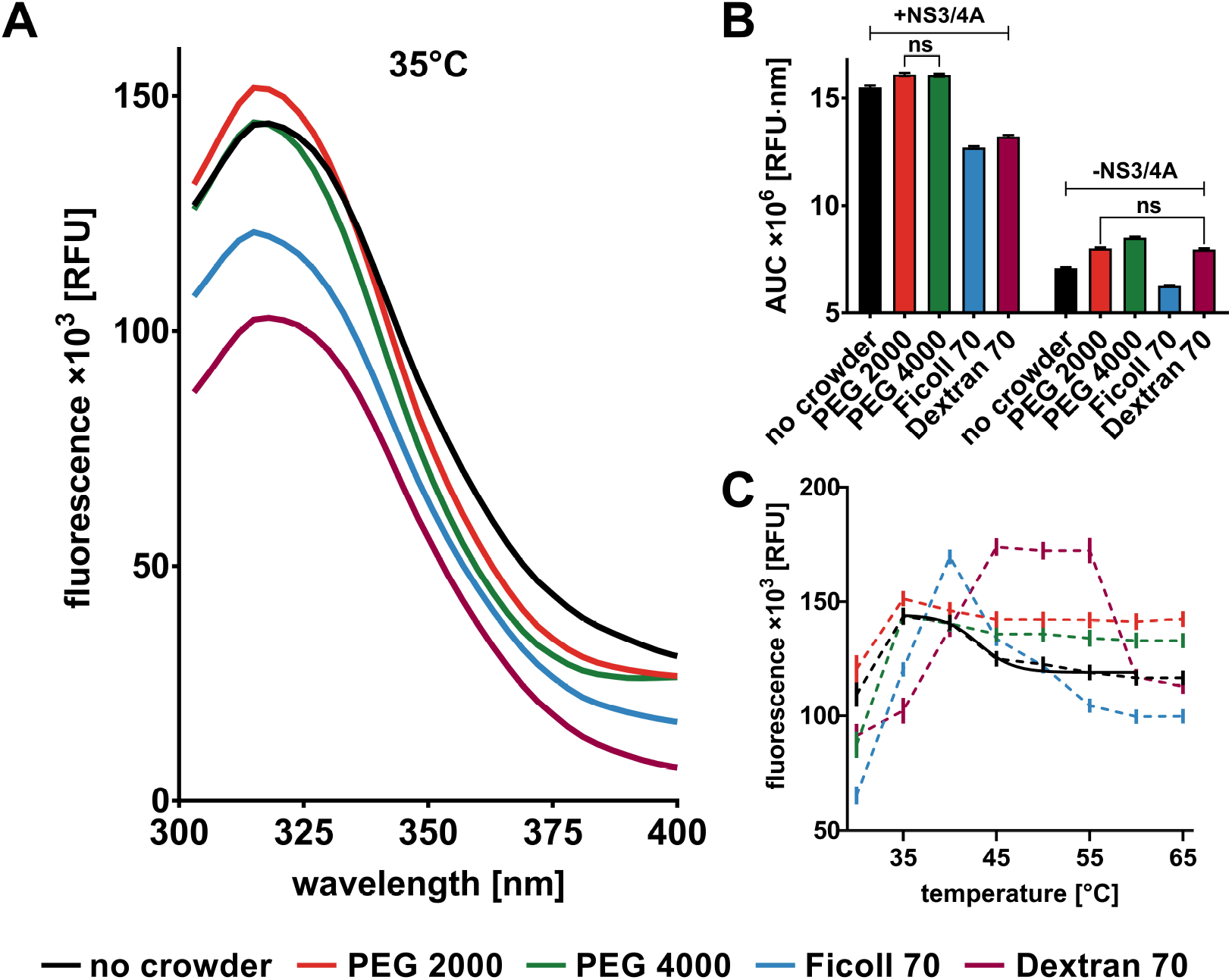
NS3/4A intrinsic fluorescence under dilute and crowded conditions. (A) Fluorescence emission spectra at 35°C were recorded with excitation at 280 nm for 1 μg/mL enzyme in dilute conditions, and in 200 g/L PEGs, 200 g/L Ficoll 70, and 150 g/L Dextran 70. Spectra were baseline-corrected and smoothed with the Savitzky-Golay method. (B) Fluorescence yield at 35°C quantified as the area under the emission curve (AUC; mean ± SD; *n* = 3). Statistical differences among conditions were assessed by two-way ANOVA and were significant (*p* < 0.0001), unless stated otherwise (not significant, ns). (C) Baseline-corrected mean fluorescence intensity (mean ± SEM; *n* = 3) as a function of temperature. For dilute conditions, data were fitted to a Boltzmann sigmoidal curve across 35-60°C, with the midpoint estimated as 43.1°C.

After adding 200 g/L of PEG 2000, the emission maximum red-shifted by 3 nm, while the overall intensity remained similar relative to dilute conditions, indicating a slightly more polar Trp environment without major quenching (**Figure 4A**). In PEG 4000, the peak position did not change, but fluorescence yield decreased by 10% (**Figure 4B**). In both PEGs, the long-wavelength tail contracted (**Figure 4A**), giving a narrower spectrum and suggesting reduced microenvironmental heterogeneity, often linked to restricted local flexibility [73]. Size-dependent effects of PEG on fluorescence have been reported in other proteins [48, 74]. Previous MD simulations showed broadly distributed PEG contacts with hydrophobic, aromatic, and sulfur-containing residues of NS3/4A [32], which may perturb local packing around Trp53 and Trp85. The subtle polarity increase in PEG 2000 likely reflects its shorter, more flexible chains accessing partially exposed hydrophobic pockets more effectively [75]. In contrast, the longer chains of PEG 4000 potentially limit penetration into these pockets, leading to different interaction patterns, for example, by displaying fewer ether oxygens for interactions [76]. Since PEG also reduced NS3/4A activity, these fluorescence changes may reflect an altered structure near the catalytically relevant regions. For instance, Trp53 contacts both NS4A and the loop that positions His57 within the catalytic triad (**Figure 1B**). Even though in our MD simulations of NS3/4A surrounded by short PEG molecules, we did not observe PEG close contacts or interactions with the active site residues [32, 34], PEG 4000 interactions with other NS3/4A residues may propagate to indirectly alter the dynamics of the active site.

In 200 g/L Ficoll 70, the NS3/4A fluorescence decreased by 20% in peak intensity and 30% in overall yield, while the emission maximum remained unchanged relative to dilute conditions (**Figure 4A-B**). The contraction of the long-wavelength tail indicates a more uniform Trp microenvironment and reduced local heterogeneity [73]. However, inert crowding is typically expected to favor compaction and reduce solvent exposure of tryptophans, leading to increased fluorescence, as reported for the cellular retinoic acid-binding protein [52]. In contrast, the observed decrease in fluorescence intensity without a major change in polarity points to enhanced non-radiative quenching, often by ligands or neighboring residues [77, 78]. Previous MD simulations indicate that ficoll interacts more extensively with NS3/4A than PEG, forming numerous hydrogen bonds with the protein surface [32]. These simulations also showed that ficoll-mimicking crowders altered NS3/4A conformational flexibility to a greater extent than PEG. The branched polysucrose structure and flexibility of ficoll in ionic solution [79, 80] likely facilitate contacts with the protein, which may promote static quenching of Trp residues through proximity to Met residues or ficoll hydroxyls, as reported for fibrinogen [81]. Despite reduced fluorescence, the NS3/4A activity increased in the presence of ficoll, indicating that the protease remained functionally folded [19]. These findings suggest that ficoll may perturb local environments without inducing global unfolding, consistent with the studies on the α-subunit of tryptophan synthase, where Ficoll 70 induced local loosening while the overall fold remained intact [53]. Fluorescence changes near Trp53, which lies at the interface critical for NS4A binding and catalytic alignment, point to localized rearrangements that coincide with increased enzymatic efficiency in ficoll.

In 150 g/L Dextran 70, NS3/4A fluorescence was strongly reduced: the maximum intensity decreased by 40%, the overall fluorescence yield dropped by 60%, and the emission maximum red-shifted by 3 nm relative to dilute conditions (**Figure 4A–B**). The long-wavelength tail contracted more than in any other crowder, indicating strongly reduced microenvironmental heterogeneity around Trp residues [73]. The combination of marked intensity loss with only a modest red-shift suggests that Trp53 and Trp85 become slightly more solvent-exposed while also experiencing enhanced non-radiative quenching. This subtle difference in local perturbation between the two polysaccharide crowders is consistent with previous reports of divergent effects on proteins such as apo-myoglobin [64], and with the distinct influence of Dextran 70 versus Ficoll 70 on NS3/4A kinetics.

These results reveal distinct local structural perturbations in NS3/4A under different crowding conditions. PEGs caused modest polarity shifts and reduced spectral heterogeneity, consistent with restricted local flexibility. In contrast, Ficoll 70 and Dextran 70 produced stronger fluorescence quenching; results with ficoll suggested enhanced non-radiative interactions, while those with dextran indicated greater polarity changes, both pointing to crowder-specific rearrangements near buried Trp residues without global unfolding.

### 2.3 NS3/4A conformational response near Trp residues to moderate heating under crowding

We further assessed local thermal unfolding profiles using temperature-dependent fluorescence. Increasing temperature from 30 to 65°C progressively altered the fluorescence spectra under all conditions (**Figure S4A**), indicating structural changes around the Trp residues. Across this range, the 350/330 nm fluorescence ratio remained below the threshold at which cooperative unfolding transitions can be reliably detected (< 0.67 [82]; **Figure S4B**), and changes in the ratio were not statistically significant. This further suggests that the observed spectral shifts reflect local perturbations rather than global unfolding.

From 30 to 35°C, fluorescence intensity increased by more than 25% in buffer, PEGs, and Ficoll 70 (**Figure 4C**). The emission maximum blue-shifted by 6–9 nm (**Figure S4A**), suggesting a more homogeneous and less polar local environment, consistent with tightening or ordering around Trp residues [73]. In Dextran 70, this initial intensity increase was absent, indicating no additional stabilization upon mild heating. In buffer, fluorescence remained stable up to 40°C. Above this temperature, intensity decreased by more than 10% and the emission maximum red-shifted to 324 nm, indicating progressive local solvent exposure [71]. A midpoint of approximately 43°C was estimated for these local changes (**Figure 4C**). Notably, no abrupt transition was observed up to 65°C, suggesting that global fold stability is preserved throughout this range, consistent with the buffer formulation designed to stabilize NS3/4A for in vitro assays [83]. As mild heating has been reported to trigger hydrophobicity-driven local folding primarily in natively disordered proteins [84], these observations align with our earlier hypothesis of an incompletely folded conformation after sample thawing on ice.

In 200 g/L PEG 2000 and PEG 4000, fluorescence intensity increased slightly during the initial heating step and then remained stable up to 65°C (**Figure 4C**), indicating no major temperature-induced perturbations in the local Trp environment. The emission maximum stayed between 315–318 nm across the entire temperature range (**Figure S4A**), consistent with a largely unchanged local polarity around both Trp residues [73]. Although the overall temperature profiles were similar for the two PEGs, fluorescence yield was slightly higher in PEG 2000 than in PEG 4000, suggesting subtle molecular-weight–dependent differences in how PEG interacts with the protein. PEG has previously been reported to stabilize local structural elements during heating [40, 85–87]. Together with the kinetic data, these fluorescence and thermal profiles indicate that PEG does not unfold NS3/4A, but likely restricts local flexibility. This reduction in conformational sampling may underlie the observed decrease in catalytic efficiency. These observations offer a potential structural explanation for why PEG slows catalytic steps while leaving substrate binding largely intact.

In 200 g/L Ficoll 70, fluorescence intensity increased up to 40°C and the emission maximum blue-shifted to 312 nm (**Figure 4C** and **Figure S4A**), indicating a more hydrophobic Trp environment and reduced collisional quenching [73]. This suggests that local packing around Trp residues becomes tighter upon moderate heating. Above 40°C, however, fluorescence intensity dropped sharply and the emission maximum shifted progressively to 324 nm by 50°C, reflecting increased polarity and local unfolding [73]. Beyond this point, the signal declined only slightly up to 65°C. Together, these patterns indicate that ficoll does not stabilize local structure during modest heating, but instead alters local folding in a more complex way than simply shifting the local unfolding midpoint. In fact, a complex influence of ficoll on protein folding was previously reported, for example, for the recombinant Rab2B protein, where Ficoll promoted the formation of partially unfolded states distinct from both native and fully unfolded conformations [88]. This ficoll-specific pattern contrasts with the behavior of dextran, whose linear, flexible chains engage the protein differently and produce distinct structural and kinetic consequences.

In 150 g/L Dextran 70, fluorescence intensity increased markedly between 35–45°C, remained stable up to 55°C, and then decreased sharply at 60°C (**Figure 4C**). The emission maximum stayed at 315 nm through 55°C and shifted to 324 nm at 60°C (**Figure S4A**), indicating a transition toward a more polar Trp environment [73]. This pattern suggests an initial reduction in local flexibility around tryptophans, followed by a plateau and then local unfolding at higher temperatures. Both polysaccharide crowders promoted a less polar, more shielded Trp environment during early heating, but Dextran 70 additionally maintained this locally folded state over the 45-55°C range. Dextran-induced thermal stability has been reported for other proteins [89, 90]. This extended local stability aligns with the reduced turnover observed in dextran, suggesting that stabilization of non-productive conformations may limit catalytic efficiency.

These results reveal that NS3/4A undergoes local structural changes without global unfolding up to 65°C. PEGs preserved spectral stability, consistent with restricted flexibility, while ficoll and dextran showed distinct thermal responses: Ficoll 70 triggered early destabilization, whereas Dextran 70 increased local structural integrity at lower temperatures and maintained it up to 55°C. These crowder-specific effects parallel their kinetic profiles, suggesting that local stability and flexibility modulate catalytic efficiency under crowded conditions.

## 3 Conclusions

We demonstrated that macromolecular crowding modulates HCV NS3/4A protease activity and thermal response in a crowder-specific manner. PEG impaired catalysis by reducing turnover and altering local flexibility without unfolding the protein. In contrast, ficoll enhanced catalytic efficiency and promoted local structural changes favorable for activity. Dextran and lysozyme inhibited activity through distinct mechanisms, likely reflecting differences in polymer structure and protein interactions. These functional effects paralleled changes in tryptophan fluorescence and thermal stability, highlighting that local structural dynamics, rather than global unfolding, play a key role in how this viral protease responds to crowded environments. These results align with broader effects reported for other enzymes and reveal NS3/4A-specific features that remain hidden in dilute in vitro systems. Understanding these environmentally induced states could inform the development of novel antiviral strategies targeting NS3/4A protease conformations that are prevalent in the cellular milieu.

## 4 Materials & methods

The HCV NS3/4A protease (genotype 1b, strain HC-J4) [91], polymer crowders, Fmoc (9-fluorenylmethoxycarbonyl)-protected amino acids, L-(+)-lactic acid, TentaGel S RAM resin (0.24 mmol/g loading), and standard peptide synthesis reagents were obtained from Sigma-Aldrich. Hen egg white lysozyme, dithiothreitol (DTT) and glycerol were from Carl Roth GmbH. EDANS (5-((2-aminoethyl)amino)naphthalene-1-sulfonic acid) was from AnaSpec Inc. All reagents were of analytical or reagent grade and were used without further purification. Data analysis and visualization were performed using GraphPad Prism v8.0.

### 4.1 FRET-substrate synthesis and purification

The NS3/4A fluorogenic substrate Ac-DED(EDANS)EEAbu-ψ-[COO]-ASK(DABCYL)-NH^2^ [92] was used (**Figure S5A**). This substrate contains a Förster resonance energy transfer (FRET) pair, in which the fluorophore EDANS is quenched by DABCYL (4-((4-(dimethylamino)phenyl)azo)benzoic acid). Enzymatic cleavage separates the EDANS fluorophore and DABCYL quencher, reducing FRET efficiency, and thereby restoring fluorescence. The FRET-substrate was synthesized manually on a 120 µmol scale using the Fmoc-based solid-phase peptide synthesis (SPPS) on Rink amide resin, as previously described [34]. After cleavage and side-chain deprotection in a trifluoroacetic acid (TFA)/water/triisopropylsilane solution (95:2.5:2.5 v/v/v, 2 h), the crude depsipeptide was purified by reversed-phase high-performance liquid chromatography (RP-HPLC) on Knauer Eurospher II 100-5 C18 columns (250 × 4.6 mm analytical; 250 × 8 mm semipreparative). A binary solvent system (S 1132, Sykam GmbH) with a 35–45% acetonitrile gradient in water containing 0.1% TFA was applied over 30 min at 1.5 mL/min (analytical) and 4.5 mL/min (semipreparative). The depsipeptide, monitored at 267 nm, eluted at 8.5 min (**Figure S5B**). The purified FRET-substrate had a purity of > 95% as determined by analytical HPLC. Identity was confirmed by electrospray ionization mass spectrometry (ESI-MS; Q-TOF Premier, Waters), yielding a mass-to-charge ratio of *m*/*z* = 1548.6, consistent with the calculated value [M + H]^+^ = 1548.6 (**Figure S5C**). TFA counterions were exchanged for chloride by dissolution in 100 mM HCl, followed by freezing and lyophilization.

### 4.2 Buffer and sample preparation

The assay buffer contained 50 mM HEPES pH 7.8, 100 mM NaCl, 20% v/v glycerol and 5 mM DTT, and was used for all dilutions. NS3/4A was diluted as needed and stored at –80°C. FRET-substrate (400 µM in assay buffer) and EDANS (1 mM in DMSO) were diluted as needed and stored at –20°C. DTT-containing solutions were thawed once and used immediately. Lysozyme was dissolved in assay buffer at room temperature, while polymer crowders were dissolved in assay buffer without DTT by mixing overnight at 35°C and 200 rpm. DTT was added fresh before enzyme incubation. NS3/4A was kept on ice, followed by 10 min preincubation at room temperature.

### 4.3 EDANS fluorescence measurement

Fluorescence was measured at 37°C in black 96-well F-bottom plates (100 µL/well; FLUOTRAC, Greiner Bio-One) using a Synergy H1 Hybrid microplate reader (BioTek) in top-read mode (7 mm read height, autogain enabled). Excitation and emission wavelengths were 340 nm and 490 nm, respectively. Plates were shaken at 37°C and 200 rpm for 30 s and allowed to settle for 30 s before the measurement. Final crowder concentrations were 50, 100, and 200 g/L for PEGs, Ficoll 70 and Dextran 70, and 15, 25, 50, and 75 g/L for lysozyme. Each condition was tested in duplicate, with samples without crowder serving as controls. Fluorescence was reported in relative fluorescence units (RFU).

#### 4.3.1 EDANS calibration

Calibration was performed in buffer and in crowder solutions using serial EDANS dilutions (0–1,000 nM final). EDANS and crowders were preincubated separately at 37°C for 10 min, then mixed in plates. Fluorescence was recorded in three independent runs, showing a linear response (R^2^ > 0.9; **Figure S6A**). To test for inner-filter effects, serial FRET-substrate dilutions (0-40 µM final) were prepared with and without 500 nM EDANS. Substrate and crowders were preincubated separately at 37°C for 10 min, then mixed in plates. Fluorescence was measured in two independent runs. As the response remained linear (R^2^ > 0.9; **Figure S6B**), no inner-filter corrections were applied in the kinetic assay.

#### 4.3.2 Enzyme kinetics assay

Kinetic measurements were performed with NS3/4A (0.05 µg/mL final, [E^tot^]) in buffer and crowder solutions, using serial FRET-substrate dilutions (0-40 µM final, [S]). Enzyme in buffer/crowder solution and substrate (in buffer only) were preincubated separately at 37°C for 10 min before mixing in the plate. Fluorescence was recorded every 45 s for 60 min in three independent runs. Additional measurements in the absence of NS3/4A were performed in duplicates under dilute conditions and in the presence of lysozyme at the highest tested concentration (75 g/L) to verify that no spontaneous FRET-substrate cleavage or HEWL-mediated hydrolysis occurred (**Figure S2**). Initial reaction velocities (*v*_0_) were determined from the linear phase of fluorescence increase and fitted to the Michaelis–Menten equation (R^2^ > 0.9, except R^2^ > 0.8 in 50 g/L lysozyme; **Figure 2A**): *v*_0_ = *v*_max_ [S]/(*K*_M_ +[S]), where *v*_max_ is maximal reaction rate and *K*_M_ is the Michaelis constant. Catalytic turnover was calculated as: *k*_cat_ = *v*_max_ /[E_tot_]. The excluded-volume fraction was estimated by approximating crowders as spheres with reported hydrodynamic radii: 1.27 nm and 1.72 nm for PEG 2000 and PEG 4000 [93], respectively, 4.70 nm for Ficoll 70 [94], 6.49 nm for Dextran 70 [95], and 1.90 nm for lysozyme [96]. The excluded volume fraction was calculated as: 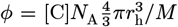, where [C] is the crowder w/v concentration, *N*_A_ is Avogadro’s number, and *M* is the crowder molecular weight [97]. The substrate concentration corrected for excluded volume effects was calculated as: [S]_eff_ = [S]/(1 − *ϕ*).

### 4.4 Intrinsic fluorescence measurements

NS3/4A (1 µg/mL final) was incubated in buffer or in 200 g/L PEGs and Ficoll 70, and 150 g/L Dextran 70 (above the overlap concentration [62]; reduced due to high viscosity below 35°C to ensure accurate handling) in sealed black 96-well F-bottom plates (100 µL/well) at room temperature for 10 min, then centrifuged at room temperature for 10 min at 2,000 rpm. Intrinsic tryptophan fluorescence was recorded on an Enspire 2300 Multilabel Reader (PerkinElmer) with excitation at 280 nm. Emission spectra were collected from 300–400 nm during a temperature ramp 30–65°C at 1°C/min, with 5°C sampling intervals and an additional 1 min equilibration delay at each step. Data for NS3/4A in dilute conditions were fitted to a Boltzmann sigmoidal model at 35-60°C (R^2^ > 0.8; **Figure 4C**). Instrument settings were: excitation/emission slit widths 5 nm, measurement height 9.5 mm, and PTM voltage 750 V. Spectra were averaged from three scans, baseline-corrected, and smoothed with the Savitzky-Golay method. The experiment was repeated twice. Fluorescence yield was quantified as the area under the emission curve (AUC) and fitted to a Boltzmann sigmoidal model for dilute conditions (R^2^ > 0.9; **Figure S4B**).

## Abbreviations

EDANS: 5-((2-aminoethyl)amino)naphthalene-1-sulfonic acid
AUC: area under the emission curve
BSA: bovine serum albumin
*k*_cat_/*K*_M_: catalytic efficiency
DABCYL: 4-((4-(dimethylamino)phenyl)azo)benzoic acid
DTT: dithiothreitol
[S] _eff_ /[S]: effective substrate concentration factor
ESI-MS: electrospray ionization mass spectrometry
ϕ: excluded volume fraction
Fmoc: 9-fluorenylmethoxycarbonyl
FRET: Förster resonance energy transfer
HEWL: hen egg white lysozyme
HCV: hepatitis C virus
*v*_0_: initial reaction rate
*v* _max_: maximal reaction rate
*K*_M_: Michaelis constant
MD: molecular dynamics
PEG: polyethylene glycol
RFU: relative fluorescence unit
RP-HPLC: reversed-phase high-performance liquid chromatography
SPPS: solid-phase peptide synthesis
TFA: trifluoroacetic acid
*k*_cat_: turnover number

## Declarations

### Data availability statement

The data are available in the supplementary material or RepOD repository at https://doi.org/10.18150/9ZPP7S

### Author contribution

ML: investigation, methodology, formal analysis, validation, writing – original draft, editing. JT: conceptualisation, project administration, funding acquisition, data curation, resources, supervision, writing – review & editing.

