## Supplementary Material for "Crowder-specific modulation of hepatitis C virus NS3/4A protease activity and local structural dynamics"

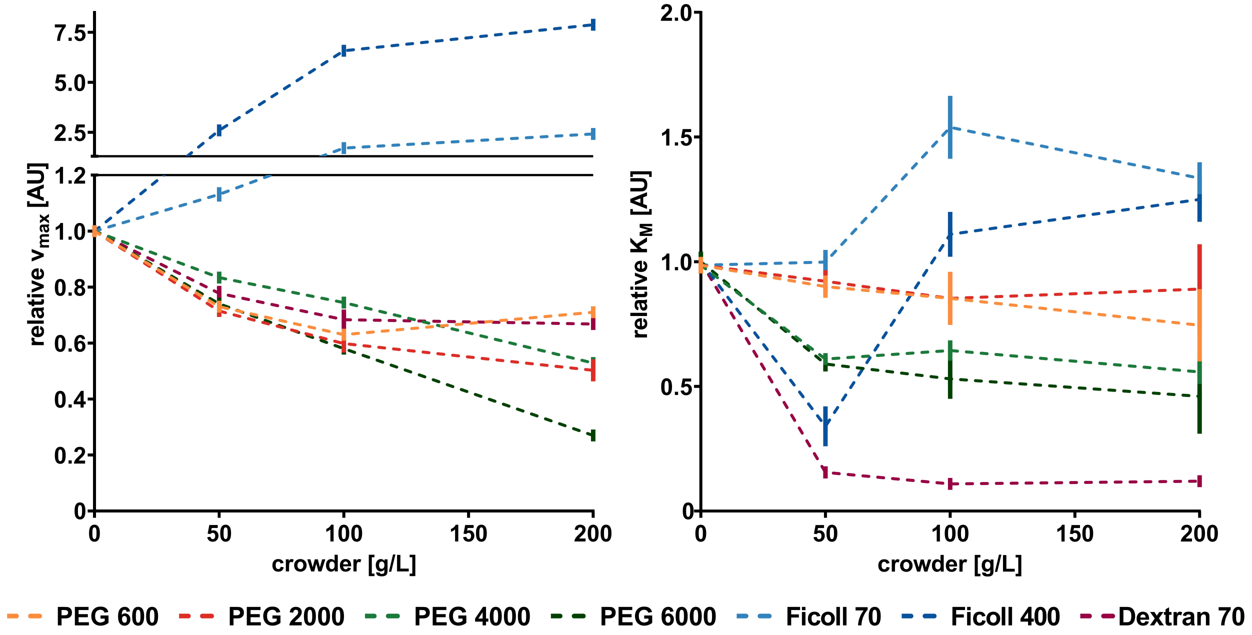


**Fig. S1 Comparison of relative kinetic parameters of the NS3/4A-mediated hydrolysis under crowded conditions.** Maximal reaction rate ($v_{\max}$) and Michaelis constant ($K_{\mathrm{M}}$) were determined from fitting initial velocities to the Michaelis-Menten equation at the 0–40 µM substrate range (**Figure 2A** in the main text) and are shown as mean ± SEM (n = 3, each in duplicate) relative to control dilute conditions. Data for PEG 600, PEG 6000, and Ficoll 400 were taken from [1].

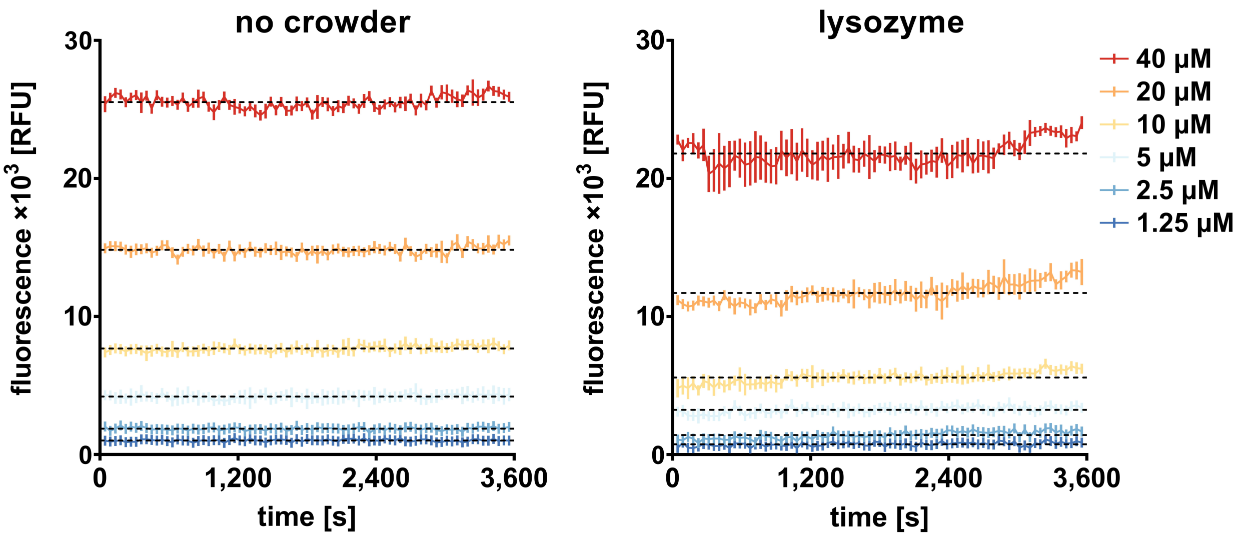


**Fig. S2 FRET-substrate fluorescence stability under dilute conditions and in the presence of lysozyme.** Fluorescence signal from 0–40 µM FRET-substrate was recorded in the absence of NS3/4A protease under dilute conditions (no crowder) and in the presence of 75 g/L lysozyme to confirm the absence of substrate cleavage. Data were baseline-corrected and are presented as mean ± SEM (n = 2). Black dashed lines represent the average fluorescence across the entire measurement period, indicating no visible increase in signal during the initial phase.

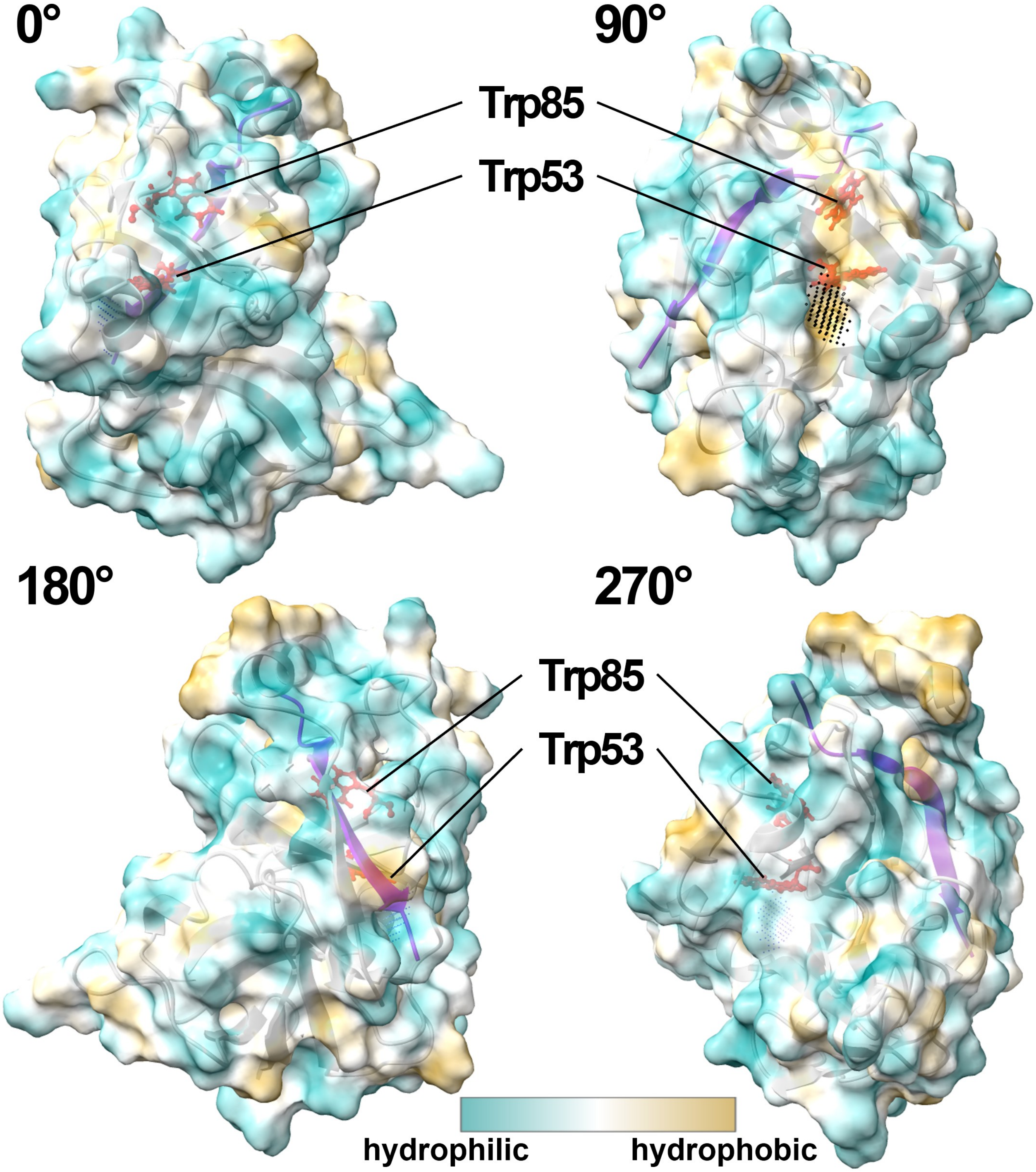


**Fig. S3 Hydrophobic surface distribution of the NS3/4A protease.** Rotational views of the NS3 protease (grey ribbon) in complex with the NS4A cofactor (violet ribbon), visualized using ChimeraX [2]. The color of the semitransparent surface maps local hydrophobicity (gold) and hydrophilicity (cyan). Tryptophans are marked in red. The solvent-exposed surface area is 4.1 Å² for Trp53, and around 1 Å² for Trp85. The hydrophobic cavity of approximately 35 Å² lined by Trp53 was identified using pyKVFinder [3] and is indicated by black dots. The inner probe radius of 1.4 Å corresponded to the approximate size of a water molecule.

**Table S1 Contacts of Trp53 in the NS3/4A protease.** Contacts were analyzed in ChimeraX [2] using the NS3/4A model based on [1] and crystal structure 4JMY [4]. Atoms were counted as contacting when located within 0.4 Å of van der Waals contact (overlap). Positive and near-zero overlap values indicate tight packing that contributes to Trp shielding. Intra-residue contacts were excluded. The chain identifier is noted as /A for the NS3 protease. Atom numbering is shown as in the PDB 4JMY structure [4].

| atom1 | atom2 | overlap [Å] | distance [Å] |
|---|---|---|---|
| /A TRP 53 HN | /A THR 46 O | 0.218 | 1.862 |
| /A TRP 53 O | /A ALA 45 HA | 0.211 | 2.269 |
| /A TRP 53 O | /A ALA 45 CA | 0.092 | 3.088 |
| /A TRP 53 O | /A THR 46 HN | 0.091 | 1.989 |
| /A TRP 53 CE2 | /A VAL 51 HG12 | 0.039 | 2.661 |
| /A TRP 53 CH2 | /A TYR 75 CD2 | 0.038 | 3.362 |
| /A TRP 53 NE1 | /A VAL 51 HB | 0.023 | 2.602 |
| /A TRP 53 CE2 | /A VAL 51 CG1 | 0.019 | 3.381 |
| /A TRP 53 HA | /A VAL 83 O | 0.002 | 2.478 |
| /A TRP 53 CE3 | /A LEU 82 HD11 | -0.035 | 2.735 |
| /A TRP 53 HZ2 | /A MET 179 SD | -0.049 | 2.831 |
| /A TRP 53 NE1 | /A VAL 51 CG1 | -0.053 | 3.378 |
| /A TRP 53 HE3 | /A LEU 82 CD1 | -0.059 | 2.759 |
| /A TRP 53 CZ2 | /A MET 175 HB1 | -0.077 | 2.777 |
| /A TRP 53 CE3 | /A LEU 82 CD1 | -0.098 | 3.498 |
| /A TRP 53 CZ2 | /A VAL 51 HG12 | -0.109 | 2.809 |
| /A TRP 53 NE1 | /A VAL 51 CB | -0.114 | 3.439 |
| /A TRP 53 HE3 | /A VAL 83 O | -0.132 | 2.612 |
| /A TRP 53 N | /A THR 46 O | -0.157 | 2.862 |
| /A TRP 53 O | /A THR 46 N | -0.174 | 2.879 |
| /A TRP 53 HE3 | /A LEU 82 HD11 | -0.189 | 2.189 |
| /A TRP 53 CA | /A VAL 83 O | -0.223 | 3.403 |
| /A TRP 53 CE3 | /A LEU 82 CG | -0.223 | 3.623 |
| /A TRP 53 HE1 | /A VAL 51 HB | -0.234 | 2.234 |
| /A TRP 53 O | /A ALA 45 CB | -0.242 | 3.422 |
| /A TRP 53 HN | /A THR 46 C | -0.244 | 2.944 |
| /A TRP 53 CE2 | /A MET 175 HB1 | -0.255 | 2.955 |
| /A TRP 53 CE3 | /A VAL 83 O | -0.265 | 3.445 |
| /A TRP 53 O | /A ALA 45 C | -0.265 | 3.445 |
| /A TRP 53 CD1 | /A VAL 48 HB | -0.277 | 2.977 |
| /A TRP 53 CD1 | /A MET 175 SD | -0.280 | 3.762 |
| /A TRP 53 HH2 | /A MET 179 HG2 | -0.290 | 2.290 |
| /A TRP 53 CD1 | /A VAL 51 O | -0.292 | 3.472 |
| /A TRP 53 CZ3 | /A LEU 82 HG | -0.295 | 2.995 |
| /A TRP 53 CH2 | /A TYR 75 CG | -0.300 | 3.700 |
| /A TRP 53 CE3 | /A LEU 82 HG | -0.301 | 3.001 |
| /A TRP 53 CZ3 | /A TYR 75 HB2 | -0.317 | 3.017 |
| /A TRP 53 HH2 | /A TYR 75 CD2 | -0.323 | 3.023 |
| /A TRP 53 CD1 | /A VAL 51 CG1 | -0.331 | 3.731 |
| /A TRP 53 CD2 | /A VAL 51 CG1 | -0.345 | 3.745 |
| /A TRP 53 HE3 | /A LEU 82 CG | -0.347 | 3.047 |
| /A TRP 53 CZ2 | /A MET 179 SD | -0.353 | 3.835 |
| /A TRP 53 O | /A THR 54 HB | -0.360 | 2.840 |
| /A TRP 53 HA | /A GLY 84 HA2 | -0.363 | 2.363 |
| /A TRP 53 CZ3 | /A LEU 82 CG | -0.374 | 3.774 |
| /A TRP 53 HH2 | /A TYR 75 CG | -0.378 | 3.078 |
| /A TRP 53 CG | /A MET 175 SD | -0.390 | 3.872 |
| /A TRP 53 HD1 | /A VAL 48 HB | -0.393 | 2.393 |
| /A TRP 53 CE3 | /A VAL 83 C | -0.394 | 3.794 |
| /A TRP 53 CD1 | /A VAL 51 HG11 | -0.395 | 3.095 |
| /A TRP 53 CZ2 | /A VAL 51 CG1 | -0.397 | 3.797 |

**Table S2 Contacts of Trp85 in the NS3/4A protease.** Contacts were analyzed as in **Table S1**. The chain identifiers are noted as /A for the NS3 protease and /D for the NS4A cofactor.

| atom1 | atom2 | overlap [Å] | distance [Å] |
|---|---|---|---|
| /A TRP 85 O | /A CYS 52 HN | 0.261 | 1.819 |
| /A TRP 85 CE2 | /A LEU 64 HB2 | 0.101 | 2.599 |
| /A TRP 85 NE1 | /A GLN 73 CG | 0.072 | 3.253 |
| /A TRP 85 HN | /A CYS 52 O | 0.024 | 2.056 |
| /A TRP 85 CD2 | /A LEU 64 HB2 | -0.009 | 2.709 |
| /A TRP 85 CD1 | /A ILE 71 O | -0.010 | 3.190 |
| /A TRP 85 CZ3 | /A THR 54 CG2 | -0.023 | 3.423 |
| /A TRP 85 HD1 | /A THR 72 C | -0.038 | 2.738 |
| /A TRP 85 O | /A VAL 51 HG13 | -0.054 | 2.534 |
| /A TRP 85 O | /A CYS 52 N | -0.068 | 2.773 |
| /A TRP 85 HD1 | /A ILE 71 O | -0.073 | 2.553 |
| /A TRP 85 NE1 | /A GLN 73 HG2 | -0.079 | 2.704 |
| /A TRP 85 HE1 | /A LYS 62 O | -0.083 | 2.163 |
| /A TRP 85 NE1 | /A LEU 64 N | -0.105 | 3.355 |
| /A TRP 85 O | /A VAL 51 HA | -0.124 | 2.604 |
| /A TRP 85 CD1 | /A GLN 73 HA | -0.137 | 2.837 |
| /A TRP 85 HA | /A PRO 86 HD1 | -0.155 | 2.155 |
| /A TRP 85 HH2 | /A ALA 59 CB | -0.165 | 2.865 |
| /A TRP 85 HZ3 | /A THR 54 CG2 | -0.176 | 2.876 |
| /A TRP 85 CD1 | /A GLN 73 N | -0.177 | 3.502 |
| /A TRP 85 CZ2 | /D VAL 225 HG21 | -0.177 | 2.877 |
| /A TRP 85 CH2 | /A ALA 59 CB | -0.181 | 3.581 |
| /A TRP 85 CD1 | /A GLN 73 CA | -0.191 | 3.591 |
| /A TRP 85 HD1 | /A GLN 73 N | -0.191 | 2.816 |
| /A TRP 85 CH2 | /A THR 54 CG2 | -0.201 | 3.601 |
| /A TRP 85 CH2 | /A THR 54 HG22 | -0.206 | 2.906 |
| /A TRP 85 O | /A VAL 51 CG1 | -0.209 | 3.389 |
| /A TRP 85 CZ3 | /32 TIP 21098 H1 | -0.213 | 2.913 |
| /A TRP 85 HH2 | /A ALA 59 HB3 | -0.213 | 2.213 |
| /A TRP 85 HB1 | /A ILE 71 CG2 | -0.218 | 2.918 |
| /A TRP 85 NE1 | /A GLN 73 HG1 | -0.218 | 2.843 |
| /A TRP 85 CZ3 | /A THR 54 HG22 | -0.225 | 2.925 |
| /A TRP 85 HZ3 | /32 TIP 21098 H1 | -0.228 | 2.228 |
| /A TRP 85 CZ3 | /A THR 54 HG23 | -0.228 | 2.928 |
| /A TRP 85 HD1 | /A THR 72 O | -0.232 | 2.712 |
| /A TRP 85 O | /A VAL 51 CA | -0.235 | 3.415 |
| /A TRP 85 CE2 | /A LEU 64 CB | -0.237 | 3.637 |
| /A TRP 85 NE1 | /A LEU 64 HB2 | -0.254 | 2.879 |
| /A TRP 85 HZ2 | /D VAL 225 HG21 | -0.255 | 2.255 |
| /A TRP 85 CD1 | /A GLN 73 CG | -0.273 | 3.673 |
| /A TRP 85 CD2 | /A LEU 64 CB | -0.281 | 3.681 |
| /A TRP 85 HZ2 | /A LYS 62 O | -0.283 | 2.763 |
| /A TRP 85 CZ2 | /A ALA 59 HB1 | -0.295 | 2.995 |
| /A TRP 85 N | /A CYS 52 O | -0.306 | 3.011 |
| /A TRP 85 CE2 | /A GLN 73 HG2 | -0.310 | 3.010 |
| /A TRP 85 CD1 | /A THR 72 C | -0.310 | 3.710 |
| /A TRP 85 CB | /A PRO 86 HD1 | -0.315 | 3.015 |
| /A TRP 85 NE1 | /A THR 63 C | -0.320 | 3.645 |
| /A TRP 85 HE3 | /A CYS 52 O | -0.321 | 2.801 |
| /A TRP 85 HE1 | /A GLN 73 CG | -0.332 | 3.032 |
| /A TRP 85 CD1 | /A LEU 64 CB | -0.334 | 3.734 |
| /A TRP 85 NE1 | /A LEU 64 CB | -0.336 | 3.661 |
| /A TRP 85 HE1 | /A GLN 73 HG1 | -0.336 | 2.336 |
| /A TRP 85 C | /A CYS 52 HN | -0.340 | 3.040 |
| /A TRP 85 CZ2 | /A LYS 62 O | -0.341 | 3.521 |
| /A TRP 85 CE3 | /A LEU 64 HD22 | -0.346 | 3.046 |
| /A TRP 85 O | /A VAL 51 C | -0.347 | 3.527 |
| /A TRP 85 NE1 | /A LYS 62 O | -0.348 | 3.053 |
| /A TRP 85 CD1 | /A GLN 73 HG2 | -0.352 | 3.052 |
| /A TRP 85 CZ2 | /D VAL 225 CG2 | -0.352 | 3.752 |
| /A TRP 85 CH2 | /D VAL 225 HG21 | -0.355 | 3.055 |
| /A TRP 85 HH2 | /A THR 54 HG22 | -0.357 | 2.357 |
| /A TRP 85 C | /A VAL 51 HG13 | -0.360 | 3.060 |
| /A TRP 85 CG | /A LEU 64 CB | -0.362 | 3.762 |
| /A TRP 85 HB1 | /A ILE 71 HG21 | -0.371 | 2.371 |
| /A TRP 85 CH2 | /A ALA 59 HB1 | -0.382 | 3.082 |
| /A TRP 85 CG | /A LEU 64 HB2 | -0.383 | 3.083 |
| /A TRP 85 HZ2 | /D VAL 225 CG2 | -0.388 | 3.088 |
| /A TRP 85 HZ3 | /A THR 54 HG22 | -0.391 | 2.391 |

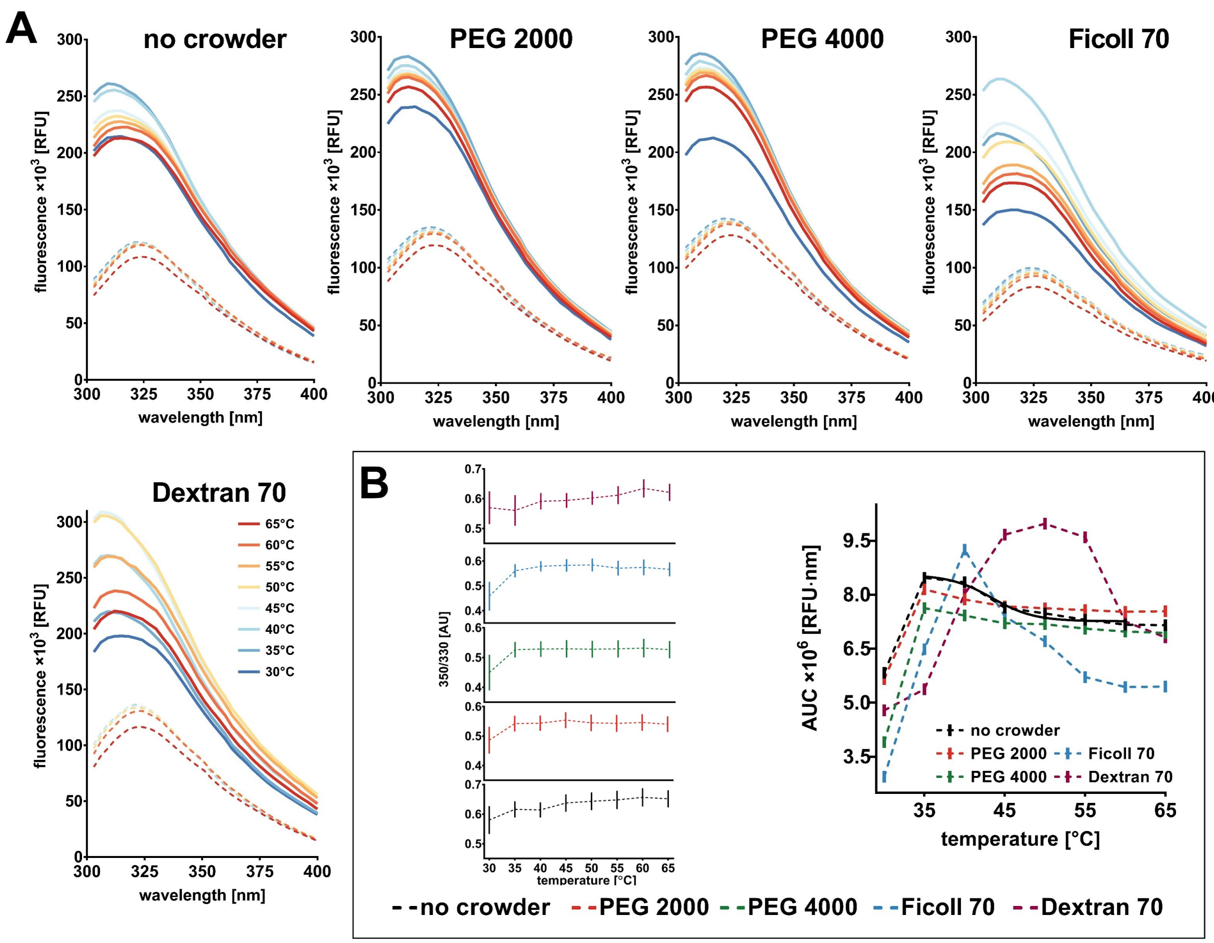


**Fig. S4 Intrinsic fluorescence of the NS3/4A enzyme under dilute and crowded conditions.** (A) Fluorescence emission spectra were recorded in control dilute conditions and in 200 g/L PEGs and Ficoll 70, and 150 g/L Dextran 70. Spectra (mean; n = 3) for protein-containing samples (solid lines) and baseline (dashed lines) were smoothed with the Savitzky-Golay method. (B) Baseline-corrected protein mean fluorescence yield quantified as the area under the emission curve (AUC; mean ± SEM) as a function of temperature. For dilute conditions, data were fitted to a Boltzmann sigmoidal curve across 35-60°C, with the midpoint estimated as 43.4°C.

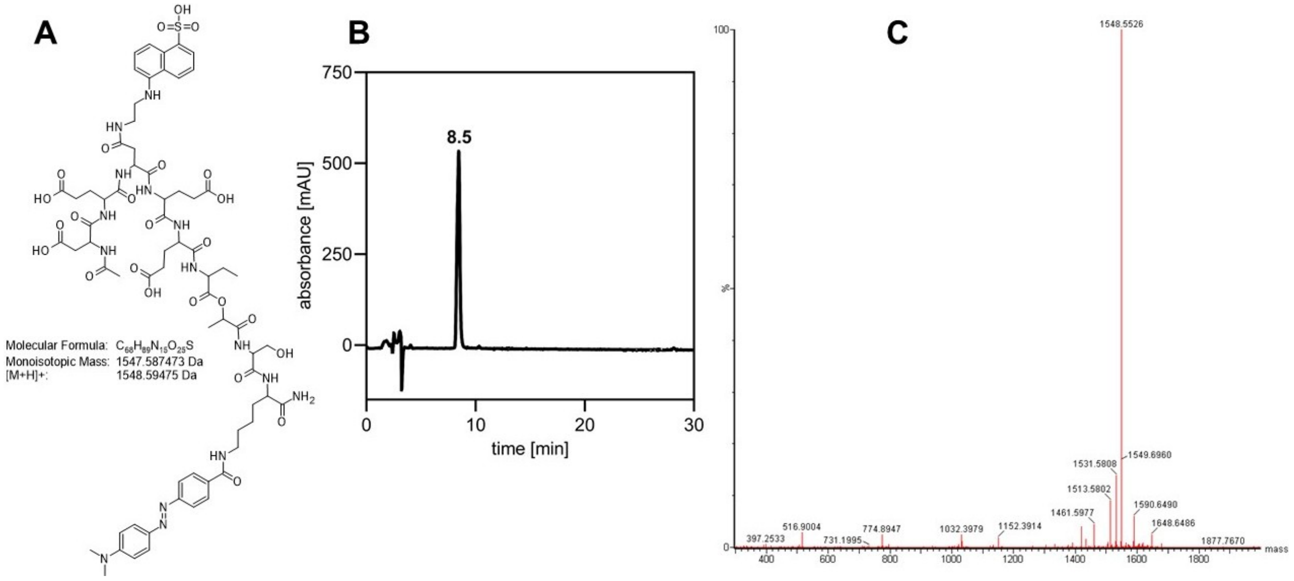


**Fig. S5 The structure, chromatogram and mass spectrum of the FRET-substrate.** (A) Chemical structure of Ac-DED(EDANS)EEAbu-ψ-[COO]-ASK(DABCYL)-$NH_2$ with calculated monoisotopic and quasi-molecular ion masses. (B) Analytical RP-HPLC chromatogram. The substrate eluted at 8.5 min. Substrate purity was determined to be $> 95\%$ by HPLC. (C) ESI-MS spectrum. The observed $m/z = 1548.6$ matched the calculated value for $[\mathrm{M + H}]^+$ confirming the product's identity.

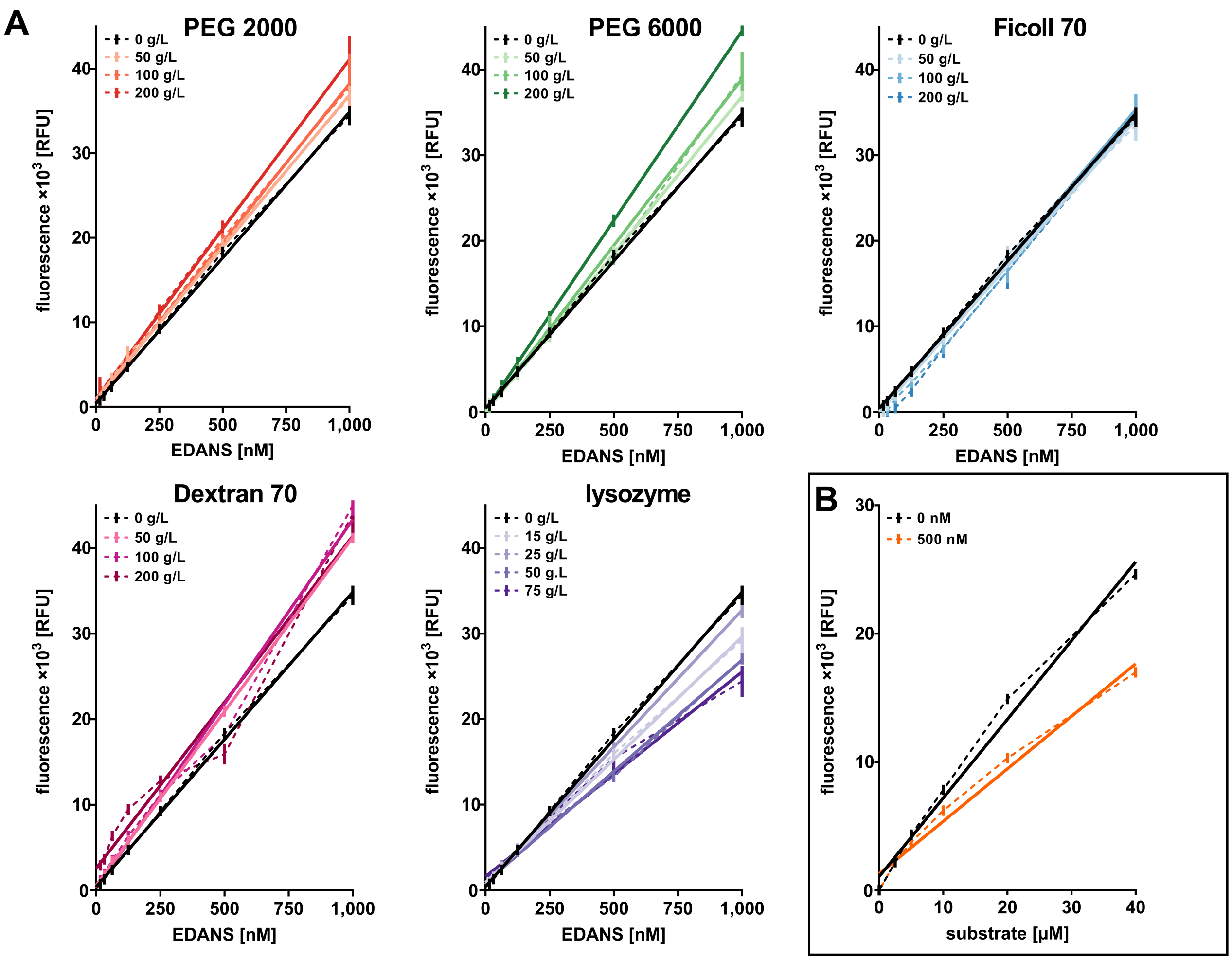


**Fig. S6 EDANS calibration under dilute and crowded conditions.** (A) Fluorescence response was tested with excitation at 340 nm and emission at 490 nm for 0-1,000 nM EDANS under dilute conditions and in 50-200 g/L polymer crowders, and 15-75 g/L lysozyme. Data points represent the mean ± SEM (n = 3, each in duplicate) and were fitted to a linear regression model (solid lines). (B) Inner filter effects were evaluated in assay buffer for 0-40 µM substrate in the absence and presence of 500 nM EDANS. Data points represent the mean ± SEM (n = 3, each in duplicate) and were fitted to a linear regression model (solid lines).
